# Place field precision during an episode predicts place field fate across episodes

**DOI:** 10.1101/2023.06.18.545503

**Authors:** YuHung Chiu, Can Dong, Seetha Krishnan, Mark E.J. Sheffield

## Abstract

Spatial memories are represented by hippocampal place cells during navigation. This spatial code is dynamic, undergoing changes across time – known as drift – and across changes in internal state, even while navigating the same spatial environment with consistent behavior. A dynamic spatial code may be a way for the hippocampus to track distinct episodes that occur at different times or during different internal states and update spatial memories. Changes to the spatial code include place fields that remap to new locations and place fields that vanish, while others are stable. However, what determines place field fate across episodes remains unclear. We measured the lap-by-lap properties of place cells in mice during navigation for a block of trials in a rewarded virtual environment. We then had mice navigate the same spatial environment for another block of trials either separated by a day (a distinct temporal episode) or during the same session but with reward removed to change reward expectation (a distinct internal state episode). We found that, as a population, place cells with remapped place fields across episodes had lower spatial precision during navigation in the initial episode. Place cells with stable or vanished place fields generally had higher spatial precision. We conclude that place cells with less precise place fields have greater spatial flexibility, allowing them to respond to, and track, distinct episodes in the same spatial environment, while place cells with precise place fields generally preserve spatial information when their fields reappear.

## Introduction

The hippocampus is known to play a role in encoding, consolidating, updating and retrieving episodic memories (***Andersen et al., 2006***). Within the hippocampal cell population, there are subsets of cells known as place cells, which exhibit spatial activity patterns corresponding to the animal’s location within a specific environment (***O’Keefe and Dostrovsky, 1971***). These locations are referred to as place fields (PFs), and as a population they provide a spatial representation of a given environment. The faithful reinstatement of hippocampal representations is thought to support memory retrieval (***Liu et al., 2012*; *Frankland et al., 2019*; *Gelbard-Sagiv et al., 2008; Josselyn et al., 2015; O’Keefe J, 1978***). However, recent findings show that spatial representations change with time and experience even when animals are navigating the same environment (***Driscoll et al., 2022; Lee et al., 2020; Hainmueller and Bartos, 2018; Keinath et al., 2022; Dong et al., 2021; Mau et al., 2020***). This phenomenon is known as representational drift and can occur during navigation of an environment from lap-to-lap, as demonstrated by many PFs shifting backwards (***Dong et al., 2021; Roth et al., 2012; Mehta et al., 2000***), and across repeated exposures (episodes) to the same environment on different days (***Ziv et al., 2013; Dong et al., 2021***). Representational drift may track time (***Mankin et al., 2012; Rubin et al., 2015; Mankin et al., 2015***) or amount of experience (***Khatib et al., 2023; Geva et al., 2023***). Similar changes to the spatial code are observed when animal’s undergo an internal state change during navigation, as demonstrated when attention or reward expectation is altered in an unchanging spatial environment (***Krishnan et al., 2022; Pettit et al., 2022***). At the single cell level, the fate of pre-existing place fields falls into one of 3 categories. First, place cells can remap their PFs to new locations. Second, PFs can vanish. Third, place fields can remain stable. However, what determines the fate of PFs across time or internal state changes remains unclear.

To investigate this, we reanalyzed previously published data (***Dong et al., 2021; Krishnan et al., 2022***), where 2-photon Ca^2+^ imaging was used to record the activity of large populations of pyramidal neurons in dorsal CA1 in head-fixed mice. Mice were placed on a treadmill and repeatedly traversed a virtual linear environment for water rewards. We defined cells with significant PFs during a block of trials in a single session and measured their lap-to-lap properties such as their spatial precision, firing rate variability, and backward shifting. We then determined PF fate in the same environment in either a subsequent block of trials separated by a day (a distinct temporal episode) or a subsequent block of trials during the same session but with reward removed to change reward expectation (a distinct internal state episode). Our findings reveal that remapped PFs across internal state or temporal episodes tended to have lower spatial precision during the initial block of trials, whereas stable and vanished PFs were associated with high spatial precision. This suggests that place cells with imprecise place fields generally possess greater spatial flexibility, providing a means for the hippocampus to respond to distinct episodes of experience and update spatial representations with new spatial information. Place cells with precise place fields, when they reappear, generally retain the same spatial information about the environment across episodes.

## Results

### Place fields that remap across two days in a familiar environment have lower spatial precision

To address whether place field characteristics during a single episode of navigation in an environment were associated with their fate during a second episode of the environment, head-fixed mice (n = 3) were trained to navigate a familiar VR environment (***Figure 1* A**) while the same populations of place cells were imaged in CA1 during 2 blocks of trials (distinct temporal episodes) separated by a day (***Figure 1* B**). The peak Ca^2+^ fluorescence on each lap traversal was used as a proxy for maximum spatial firing position on each lap. Peaks were then treated as events in 2D parameter space (time and spatial position). Clusters of events with consistent spatial position were identified as PFs (see Methods). The mean spatial position was then calculated and the same analysis was done the following day. PFs calculated on day 1 were then defined as either stable, remapped, or vanished based on their mean activity on day 2 (see PF categorization). Note that in this paper we distinguish between PFs that change spatial position (referred as ‘remapped’) and PFs that disappear (referred as ‘vanished’).

**Figure 1.**
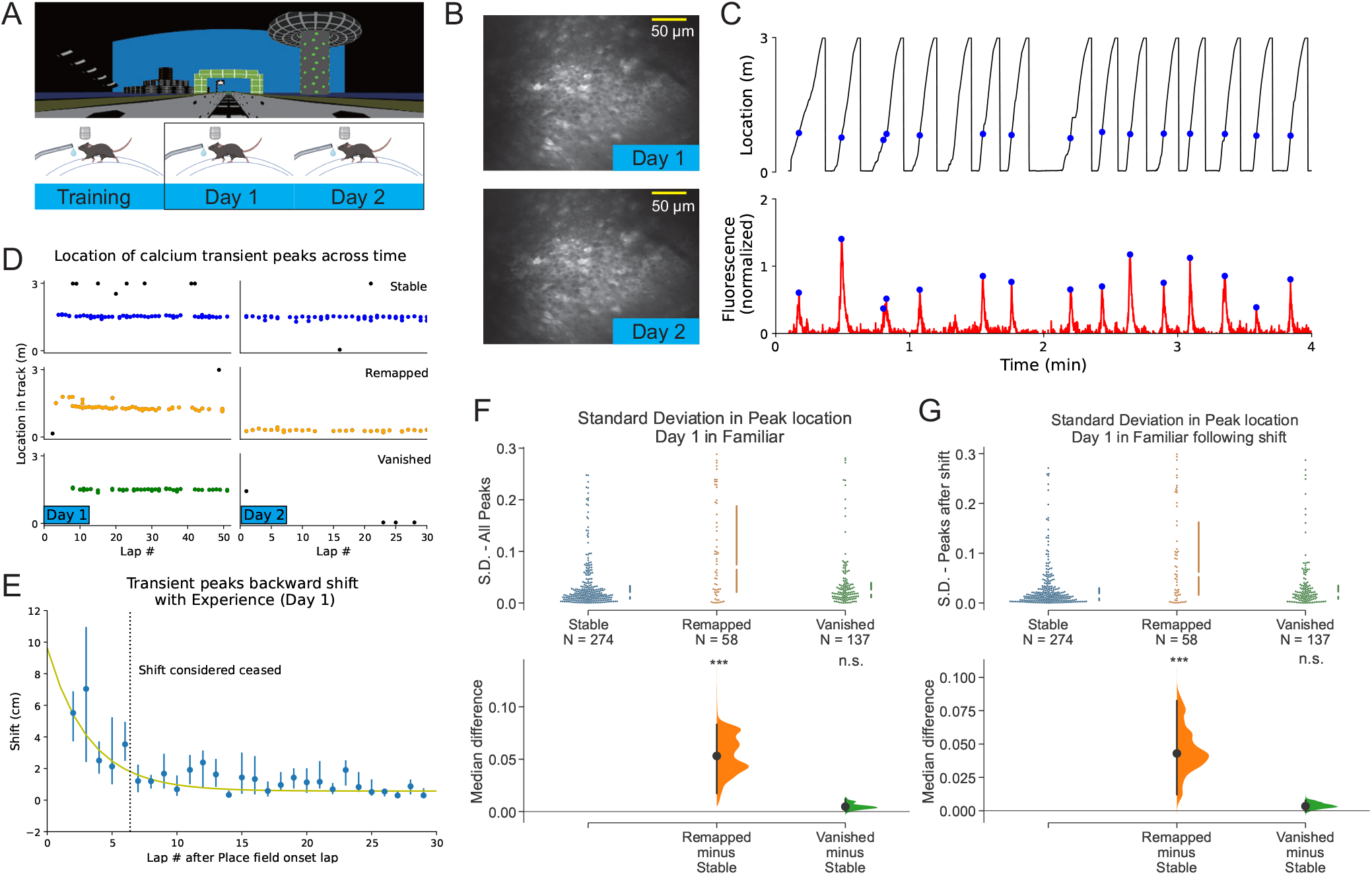
Place fields that remap across two days in a familiar environment have lower spatial precision. (A) Experimental design. Top: the familiar virtual reality (VR) environment. Bottom: Animals trained and recorded in the same VR environment. Graphic created with BioRender.com. (B) Example field of view showing imaging of the same cells on Day 1 and 2. (C) Top: Behavior of a single animal showing track location. Blue dots indicate Ca^2+^ fluorescence peaks from an example cell relative to the animal’s track location. Bottom: Fluorescence trace of the example cell across time. Blue dots indicating the peak of the Fluorescence change. (D) Examples of Stable, Remapped and Vanished place fields across days. Coloured (blue, orange and green) dots indicate in-field fluorescence peaks. Black dots are out of field peaks. (E) Average population backward shift of PF peaks on day 1. PFs are aligned to their onset lap. Line indicates fitted exponential curve: *F* (*x*) = *Ae*^−*x*/*T*^, with *A* = 9.1 ± 4.3 cm, *T* = 3.2 ± 1.1 laps (F) Comparison of spatial precision of PFs (469 PFs in day 1) from the 3 categories by measuring the standard deviation of the lap-by-lap peak locations of Stable (274/469; 58.4%), Remapped (58/469; 12.4%) and Vanished (137/469; 29.2%) PFs. Bottom: bootstrapped median difference between the three groups. 5000 re-samples, ***P < 0.001 for Remapped vs Stable, P = 0.140 for Vanished vs Stable (G) Same as (F), but includes only peaks after backward shifting. 5000 re-samples, ***P < 0.001 for Remapped vs Stable, P = 0.248 for Vanished vs Stable **Figure 1—figure supplement 1**. Following the initial episode, remapped and stable PFs have similar spatial precision during subsequent episodes **Figure 1—figure supplement 2**. Other place field metrics are not associated with place field fate across days in a familiar environment

Various properties of individual PFs in each category were then analyzed on day 1. First, lap-wise spatial precision was quantified using the standard deviation (SD) of spatial locations of the fluorescent peaks. We compared the spatial precision of PFs across the 3 categories (***Figure 1* F**). The PFs that remapped on day 2 exhibited a significantly lower spatial precision on day 1, with a higher median SD than stable PFs (P < 0.001). Vanished PFs showed no significant difference compared to stable PFs.

Studies have reported a type of drift on a lap-by-lap basis that occurs during navigation - also known as PF backward shifting (***Dong et al., 2021; Khatib et al., 2023; Geva et al., 2023; Mehta et al., 2000; Lee and Knierim, 2007; Roth et al., 2012***). Backward shifting could reduce the lap-wise precision of PFs as we measured it here. To determine whether backward shifting contributed to our measure of precision and its association with PF fate, we measured the extent of backward shifting on day 1 (***Figure 1 E***). Further, because not all PFs emerge immediately when mice start navigating a familiar environment on any particular day, we first defined the PF onset lap for each PF (***Sheffield et al., 2017; Dong et al., 2021***). Aligning PFs to their onset lap, we found that backward shifting ceased after a finite number of laps, and the decay of shifting could be well fitted to an exponential (***Figure 1 E***). We estimated the time constant *T* of the decay. We considered PFs to have ceased backward shifting after 2*T* laps from their onset, as this is the point at which 90% of the shifting had decayed (see PF backward shifting). Then, the same comparison was performed as ***Figure 1 F***, but restricted to PF activity following the backward shifting. We found that the association between PF precision and PF fate across days was maintained even when backward shifting on day 1 was excluded from the analysis ***Figure 1* G**.

We next checked if the remapped and stable PFs continued to have differences in spatial precision on day 2 (***Figure 1—figure Supplement 1 A***). We found no such difference, showing that on day 2, remapped and stable PFs have the same median precision. While the spatial precision on day 1 is relevant to PF fate across days, we next tested if other PF properties on day 1 were associated with PF fate. We first asked if the extent of backward shifting of PFs was associated with their fate. In ***Figure 1—figure Supplement 1 A***, we show the lap-by-lap shifting of all the PFs, and separately, the PFs from each category fitted to an exponential. Remapped, stable, and vanished PFs showed similar backward shifting dynamics. Comparing both the amplitudes and the time constants for the exponential fits along with their uncertainty values for all the categories demonstrated no differences between the groups (***Figure 1—figure Supplement 1A).***

Next, we asked if PF firing rates were associated with PF fate. Using peak amplitudes of calcium transients as a proxy for max firing rate on each PF traversal, we quantified the deviation in amplitudes from lap-to-lap. Comparing this measure between the 3 categories of PF fate revealed no significant difference (***Figure 1—figure Supplement 2 B***).

Not only do PFs emerge on different laps in a familiar environment, place cells can stop firing in their PF before the session ends. The PF onset lap and PF end lap, as well as the total laps in between onset and end (PF duration), can therefore be quantified for each PF. ***Figure 1—figure Supplement 2 C*** shows the histograms of PF onset laps, end laps and total laps for the three PF fate categories. We found no differences between the PF fate categories.

Together, our investigation into place field properties and PF fate across days in a familiar environment suggests that it is randomly varying lap-by-lap spatial dynamics on day 1 that is related to the across-day fate of PFs, and other PF properties are unrelated.

### Place fields that remap across two days in a novel environment have lower spatial precision

When mice are introduced to a novel environment, global remapping occurs in CA1 in which a new map forms (***Colgin et al., 2008; Sheffield et al., 2017; Dong et al., 2021***). Once the PFs that comprise the new map emerge, they typically are less precise than in familiar environments (***Frank et al., 2004***). We therefore tested if the relationship between lap-wise precision and across-day PF fate that we observed in a familiar environment also occurred in a novel environment during familiarization. We therefore switched mice (n = 3) to a novel VR environment while imaging CA1 (***Figure 2 A-B***) and identified PFs (***Figure 2 C***). We first wanted to determine how the newly-formed PF map backward shifted from lap-to-lap on day 1 (Fig. 2D). We found backward shifting was prolonged compared to the familiar environment (*T* = 3.2 ± 1.1 laps for familiar environment and *T* = 5.3 ± 1.0 laps for novel environment) as previously reported (***Dong et al., 2021***). We then measured PF precision on day 1 and compared between the three categories of PFs based on their fate on day 2. Once again, we observed that remapped PFs had lower spatial precision than stable and vanished PFs (Fig. 2F), with a higher median SD, even when backward shifting was excluded from the precision analysis (***Figure 2G***). Also, just as in a familiar environment, day 2 lap-by-lap precision showed no difference between remapped and stable PFs ***Figure 1—figure Supplement 1***, and no other PF properties measured on day 1 were related to PF fate on day 2 (***Figure 2—figure Supplement 2***).

**Figure 2.**
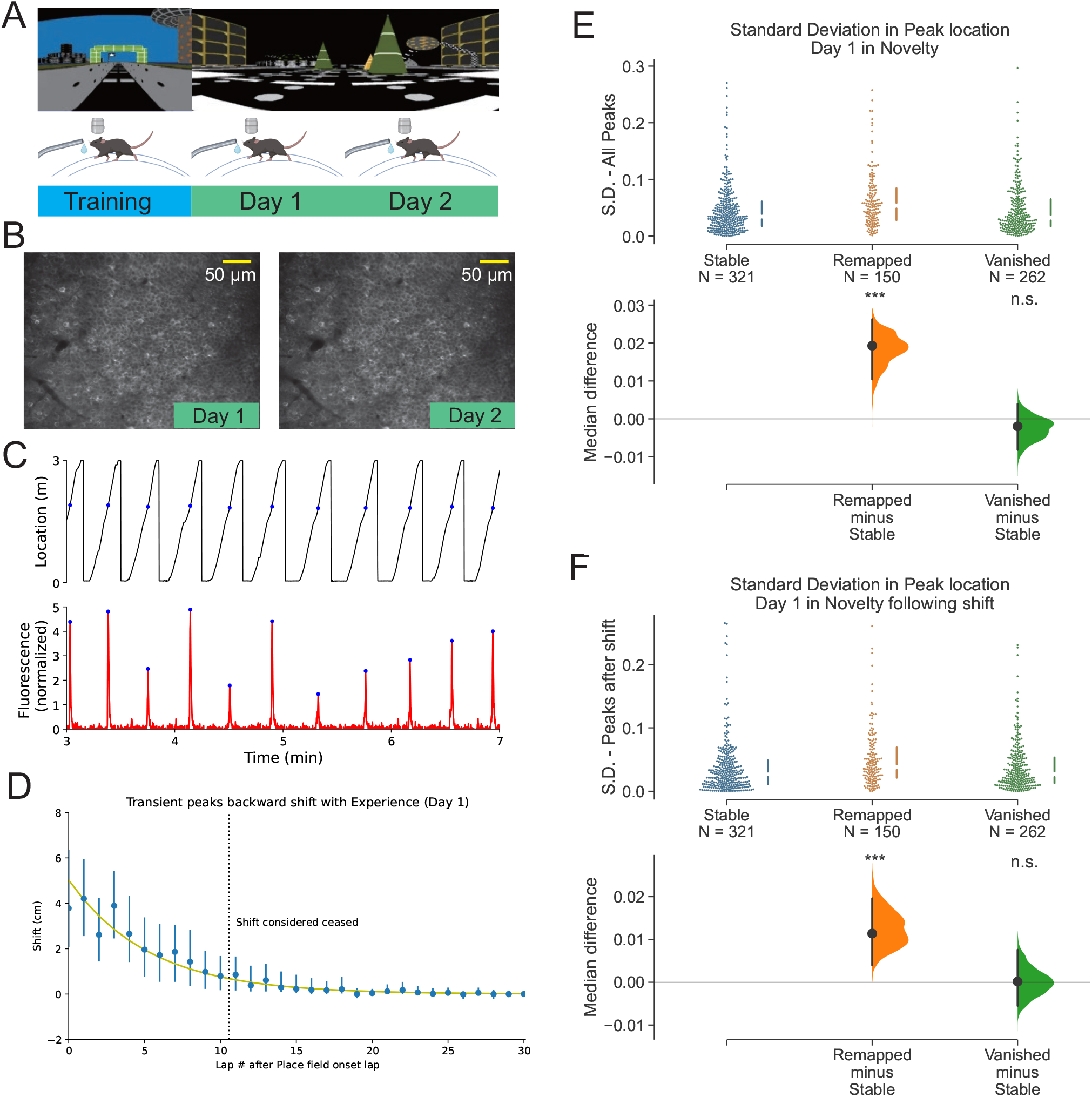
Place fields that remap across two days in a novel environment have lower spatial precision. (A) Experimental design. Top: the familiar environment (F) and the novel environment (N). Bottom: The animals were trained in a familiar environment, then switched to a novel environment and imaged for two days. Graphic created with BioRender.com. (B) Example field of view for Day 1 and 2 showing the same imaged cells. (C) Top: Behaviour of a single animal showing track location. Blue dots indicate the animals location when the example cell’s calcium transient is at its peak. Bottom: Time-series fluorescent trace for an example cell. Blue dots indicating the transient peaks. (D) Average population backward shifting of PF peaks within the session on day 1. PFs are aligned to their onset lap. Line indicates fitted exponential curve: *F* (*x*) = *Ae*^−*x*/*T*^, with *A* = 5.0 ± 1.3 cm, *T* = 5.3 ± 1.0 laps (E) Comparison of spatial precision of PFs (733 PFs on day 1) from the 3 categories by measuring the standard deviation of the lap-by-lap peak locations of Stable (321/733, 43.8%), Remapped (150/733, 20.5%) and Vanished (262/733, 35.7%) PFs. Bottom: bootstrapped median difference between the three groups. 5000 re-samples, ***P < 0.001 for Remapped vs Stable, P = 0.383 for Vanished vs Stable. (F) Same as (E), but includes only peaks after the backward shifting. 5000 re-samples, ***P < 0.001 for Remapped vs Stable, P = 0.959 for Vanished vs Stable. **Figure 2—figure supplement 1**. Other place field metrics are not associated with place field fate across days in a novel environment

### Place fields that remap in response to changed reward expectation tend to have lower spatial precision

A recent study showed that some PFs remap when the internal state of reward expectation changes in an unchanging spatial environment (***Krishnan et al., 2022***). We therefore asked whether lap-by-lap spatial precision of PFs was associated with remapping under these conditions of altered internal state. To do this, mice were trained and then imaged in the same familiar rewarded environment (***Figure 3 A***). Trained mice (n = 5) were first water rewarded for a block of trials and then reward was removed for a subsequent block of trials (Unrewarded condition: UR). After a few laps, mice stopped pre-emptively licking for reward, demonstrating a loss of reward expectation (***Krishnan et al., 2022***). Then, reward was reintroduced (Re-rewarded condition: RR)

**Figure 3.**
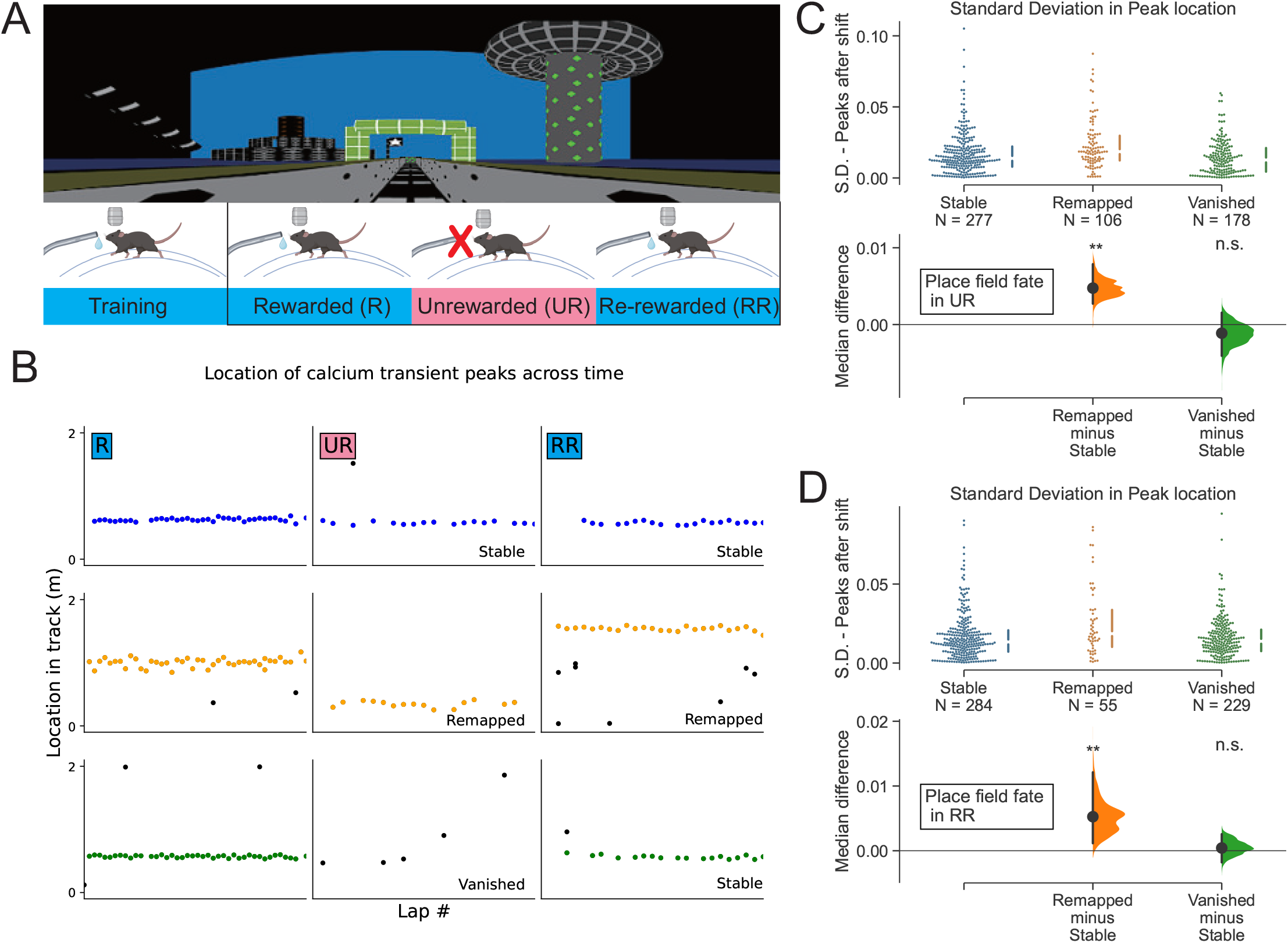
Place fields that remap in response to changed reward expectation tend to have lower spatial precision. (A) Experimental design. Top: the familiar VR environment. Bottom: The Animals were trained and recorded in the same VR environment during changes in reward. Graphic created with BioRender.com (B) Example of stable (top), remapped (middle) and vanished (bottom) place fields across changes in reward expectation. Coloured (blue, orange and green) dots indicate in-field transient peaks. The place field can undergo remapping in UR and then again in RR. (C) Comparison of spatial precision of PFs (561 PFs in R, see Defining PFs) from the 3 categories by measuring the standard deviation of the lap-by-lap peak locations of stable (277/561, 49.4%), remapped (106/561, 18.9%) and vanished (178/561, 31.7%) PFs between R and UR. Bottom: bootstrapped median difference between the three groups. 5000 re-samples, **P = 0.0016 for remapped vs stable, P = 0.480 for vanished vs stable (D) Same as (E), but comparison between R and RR. (568 PFs in R, see Defining PFs; Stable PFs: 284/568, 50.0%; Remapped PFs: 55/568, 9.7%; Vanished PFs: 229/568, 40.3%) 5000 re-samples, **P = 0.0022 for remapped vs stable, P = 0.683 for vanished vs stable **Figure 3—figure supplement 1**. Other place field metrics in R are not associated with place field fate in UR **Figure 3—figure supplement 2**. Other place field metrics in R are not associated with place field fate in RR

***Figure 3 B*** shows example place cells; one with a stable PF across the R-UR-RR conditions (top), one with a remapped PF across all conditions (middle), and one PF that vanished in UR but reappeared in RR (bottom). We then investigated whether the lap-by-lap spatial precision of PFs in R determined their fate in UR or RR (***Figure 3 C, D***). Similar to the fate of PFs across days, we found that the remapped PFs in UR (Fig. 3C) and RR (Fig. 3D) exhibited lower spatial precision in R. Again, other PF properties had no association with remapping or stability ***Figure 3—figure Supplement 1*** and ***Figure 3—figure Supplement 2***. This indicates that across the population, remapped PFs caused by internal state changes tend to have lower precision than stable/vanished PFs, behaving similarly to remapped PFs across days.

## Discussion

We investigated hippocampal CA1 spatial code dynamics occurring across episodes in unchanging spatial environments to determine whether firing characteristics during a single episode was associated with how cells encode future episodes. Our findings show that PFs that remapped across episodes separated in time by a day, or across episodes distinguished by differences in reward expectation, had a statistically lower lap-by-lap spatial precision during the initial episode, compared to stable and vanished PFs. This held true across episodes in novel environments as mice underwent familiarization (a form of learning). Other lap-by-lap characteristics of PFs, such as firing rate variability, backward shifting dynamics, PF onset, PF offset, and PF duration were not associated with the fate of PFs across episodes. This indicates that the spatial firing precision of PFs during navigation is related to their tendency to remap or stabilize/vanish across distinct episodes.

Drift across time has been observed in different parts of the brain (***Deitch et al., 2021; Marks and Goard, 2021; Driscoll et al., 2017; Schoonover et al., 2021***). In the hippocampus, an accurate representation of the spatial environment is preserved during drift (***Ziv et al., 2013; Keinath et al., 2022***), suggesting drift may encode non-spatial factors of the context such as time (***Mankin et al., 2012, 2015***), and experience (***Khatib et al., 2023; Geva et al., 2023***). Our data suggest that the dynamics of drift across episodes is related to the cellular activity within an episode. This also holds true for episodes that are separated by internal state changes. The dynamics of the hippocampal spatial code across episodes may therefore not be random and may instead be predictable. However, the extent of predictability remains to be directly tested.

A possible explanation for the relationship between lap-by-lap dynamics and the tendency to remap is that those PFs with less spatial precision simply receive higher variability in the activation of the CA3 inputs they receive (***Zutshi et al., 2022; Davoudi and Foster, 2019; Devalle and Roxin, 2022***). Alternatively, evidence suggests that all CA1 pyramidal cells may receive synaptic input regarding all locations in an environment from CA3 (***Grienberger et al., 2017***). What determines whether a cell fires at a given location may therefore be the strength of synapses activated at particular locations. Indeed, dendritic spikes in CA1 place cells, which is a reflection of strong synaptic input to a dendritic branch, is associated with PF stability across days (***Sheffield and Dombeck, 2015***). One idea is that these strong synapses may have undergone Hebbian potentiation, and together with the resultant somatic firing may induce homeostatic mechanisms that lower overall cellular excitability (through synaptic or intrinsic excitability renormalization) (***Miller, 1996***) and make other sets of synapses too weak to cause somatic firing in a winner-takes-all manner (***Sheffield and Dombeck, 2015; Barry and Burgess, 2007***). This process would result in a precise PF as the cell would fire only in response to those specific inputs. The dendritic spikes associated with this strong input may further serve to maintain synaptic strength to stabilize the PF across days (***Sheffield and Dombeck, 2015***). On the other hand, PFs with more lap-wise fluctuations in spatial firing may reflect differences in the sets of synapses activated from lap-to-lap. Such variations may not engage Hebbian potentiation and thus avoid the homeostatic winner-takes-all process described above. This would both cause the cell to be less precise from lap-to-lap but also allow the cell more flexibility to respond to new sets of synaptic activation that may occur across distinct episodes of experience, allowing for continuous encoding of new information in the hippocampus.

Our results also show that the PFs that vanished across episodes are indistinguishable from the stable ones in terms of spatial precision. This aligns with previous literature (***Ziv et al., 2013***), that place cells enter and exit an active subset of an underlying stable map. When these cells are active again, their PFs retain their locations (***Ziv et al., 2013***). The PFs that vanished across episodes may actually be part of this stable map, they are just not participating in the active subset during a particular episode. This is likely due to the CA3 inputs that could drive them to fire not being activated.

Overall, our study presents how lap-by-lap dynamics of PFs during an episode of navigation relate to spatial code dynamics across episodes in the same environment. This work provides insight into why some cells remap and others remain stable/vanish. It also provides insight into the synaptic mechanisms which may facilitate these cellular dynamics to support episodic memory encoding.

## Methods

### Subjects

All experimental and surgical procedures were in accordance with the University of Chicago Animal Care and Use Committee guidelines. For this study, 10-12-week-old male C57BL/6J wildtype (WT) mice (23-33g) were individually housed in a reverse 12 h light/dark cycle with an ambient temperature of ∼20 °C and ∼50% humidity. Male mice were used over female mice due to the size and weight of the headplates (9.1mm×31.7mm, ∼2g), which were difficult to firmly attach to smaller female skulls. All training and experiments were conducted during the animal’s dark cycle.

### Mouse surgery and virus injection

Mice were anesthetized (∼1-2% isoflurane) and injected with 0.5 mL of saline (intraperitoneal injection) and ∼0.45 mL of meloxicam (1–2 mg/kg, subcutaneous injection). For CA1 population imaging, a small (∼0.5-1.0 mm) craniotomy was made over the hippocampus CA1 (1.7mm lateral, -2.3 mm caudal of Bregma). A genetically encoded calcium indicator, AAV1-CamKII-GCaMP6f (Addgene, #100834) was injected into CA1 (∼75 nl) at a depth of 1.25 mm below the surface of the dura using a beveled glass micropipette. Afterwards, the site was covered up using dental cement (Metabond, Parkell Corporation) and a metal head-plate (9.1 mm × 31.7 mm, Atlas Tool and Die Works) was also attached to the skull with the cement. Mice were separated into individual cages and water restriction began the following day (0.8–1.0 ml per day). Around 7 days later, mice underwent another surgery to implant a hippocampal window as previously described (***Dombeck et al., 2010***). Following implantation, the head plate was reattached with the addition of a head ring cemented on top of the head plate which was used to house the microscope objective and block out ambient light. Post-surgery, mice were given 2-3ml of water/day for 3 days to enhance recovery before returning to the reduced water schedule (0.8-1.0 ml/day).

### Behaviour and Calcium imaging

We analyzed previously published data (***Dong et al., 2021; Krishnan et al., 2022***).

For the experiment across days (***Dong et al., 2021***), mice (n = 5 in total) were trained to run on a treadmill along a 3 m VR linear track with 4 um of water reward delivered at the end of the track (the familiar environment (F)). Mice (n = 3, #1, 2, 3) were then imaged over two days in the same environment. Calcium activity in CA1 pyramidal neurons (n = 1282 neurons) were extracted using customized MATLAB script (***Sheffield et al., 2017***), with parameters and procedures detailed in ***Dong et al***. (***2021***). For the experiment involving the novel environment (N) switch, mice were trained to run on a treadmill in F, and then on imaging days, the mice (n = 3, #3, 4, 5, #3 was also imaged for the experiment in familiar experiment) were introduced to N with different 3D visual cues but the same reward location and track length as F. Again, calcium activity of CA1 pyramidal neurons (n = 1704 neurons) were extracted with the same approach.

For the experiment with change in reward contingencies (***Krishnan et al., 2022***), mice were trained on a 2 m VR linear track for water reward. Well-trained mice showed pre-emptive licking before the reward location. On experimental day, the mice (n = 5) ran in the environment with reward (R), then the reward was unexpectedly removed (UR). The reward was then re-introduced (RR). Each condition (R, UR and RR) lasted 8-10 minutes. Population activity of CA1 pyramidal neurons (n = 1288) were measured with Ca^2+^ imaging, across the conditions. Calcium transients were extracted using suite2p (***Pachitariu et al., 2017***) as in ***Krishnan et al***. (***2022***).

### Defining PFs

After extracting significant calcium transients, we correlated the transients to the animals’ behaviour. We obtained the significant peak of each transient by finding the local maximum of a transient that exceeds the mean Δ*F* /*F* by 3 inter-quartile range of Δ*F* /*F* in a time window of 20 frames, to avoid including peaks from noise. The corresponding animal location on the track were then obtained. Fluorescence peaks were treated as events in a 2-D parameter space (time and location). PFs were then defined by event clusters in the 2D parameter space using the clustering algorithm DBSCAN (***Ester et al., 1996***), a density based clustering algorithm. A cluster then needed to include events from at least 10 different laps to be considered a PF.

The vast majority of cells had either a single cluster or no cluster in any given episode. In the limited number of cells that had multiple clusters in either episode, each cluster was treated as independent PF. For the very few cells with multiple clusters in episode 1 and episode 2, the clusters across the episodes were “paired”, such that clusters in episode 1 were paired with the closet ones in episode 2 and analyzed as such. In cases with two clusters in episode 1 and one cluster in episode 2, we considered the clusters to have merged across episodes If both clusters in the first episode were within 40 cm of the episode 2 cluster (if one or both were not, we did not consider them merged and treated each independently). The two clusters in episode 1 were therefore counted as one stable PF and we combined their spatial precision values (see PF properties on how we did this), to avoid over-counting the stable PFs. This caused the difference in total number of PFs in R for ***Figure 3* C** and ***Figure 3* D**, due to cells with multiple clusters in R having different fates in UR and RR.

### PF properties

Once clusters were identified as PFs, the time, location and transient peak Δ*F* /*F* for each event within the cluster were quantified. To determine the onset and offset of the PF, we defined the PF onset lap as the lap number of the first event in the cluster, and the PF offset lap as the lap number of the last event in the cluster. The duration of the PF was then calculated as the difference between the onset and offset lap.

The spatial location of the PF was defined as the median of the locations of the events within the cluster. The spatial precision of the PF was quantified using the standard deviation (SD) of the location in the cluster. To ensure that this measurement was not influenced by the edges of the track, PFs located within 10 cm of the track ends were excluded from the analysis.

In the rare cases where two PFs merged into a single PF across episodes (see Defining PFs), we combined their precision measure into a single value and only counted it as a single stable PF. To obtain a single precision value from the two PFs, we obtained the location of each event from each cluster and then subtracted the mean location of that cluster from each event 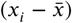. Then, the two mean-removed sets of locations were merged and the spatial precision was measured by SD of the merged set. This was performed to determine the combined spatial precision of the two PFs, that we considered a single stable PF for this analysis.

The PF firing rate dynamics were investigated by using the Δ*F* /*F* peak amplitude as a proxy for the maximum firing rate. The lap-by-lap firing rate dynamics were measured by calculating the deviation in the peak amplitude of events within the cluster.

### PF categorization

To assess the spatial stability of PFs, we categorized them based on their change in spatial location across days/conditions.

1. Stable - The PF is present on both days/conditions and any change in PF position is: Δ < 40 cm
2. Remapped - The PF is present on both days/conditions but changes PF position: Δ > 40 cm
3. Vanished - The PF is only present on the first day/condition

The specific choice of 40 cm as the threshold was made to ensure that it was larger than the typical fluctuation observed in peak locations within clusters in the dataset (a measure of PF width). This means that a change in location beyond 40 cm would typically mean a non-overlapping PF.

### PF backward shifting

PFs were aligned by their onset lap, which was defined as the lap number of the first peak in that cluster. Then the spatial positions of the PFs on each lap were obtained with a sliding window of 5 laps. The 5-lap sliding average position of individual PFs were then compared to the median location calculated from laps beyond the 15th lap from the onset lap for that PF. For each lap from the PF onset lap, the average shift over the population of PFs was calculated and plotted. An exponential fit (least square fit, scipy.optimize.curve_fit) was then applied to the trend. The error of the fitting parameters were obtained by the square-root of the diagonal elements of the covariance matrix (returned by curve_fit). The same trend was observed without the smoothing.

PFs were considered to have ceased systematically backward shifting after 2*T* laps from their onset, as the decay in shift is reduced to *e*^−2^ = 13.5% at 2*T*. Peaks after 2*T* laps were included in the comparisons that restrict to PF activity following backward shifting. Similar trends were observed using different choices than 2*T* for the cutoff.

### Statistics

For the plots regarding PF backward shifting, the error bars represent mean ± SEM.

To generate the plots comparing the spatial precision, we used the package DABEST (‘data analysis with bootstrap-coupled estimation’) (***Ho et al., 2019***). The median difference between the distributions and its confidence level were obtained with bootstrapping (5000 re-samples). P-values of the non-parametric two-sided approximate permutation t-test were reported.

## Acknowledgement

We thank D. Goodsmith and A. Madar for manuscript comments. This work was supported by: The Whitehall Foundation, The Searle Scholars Program, The Sloan Foundation, The University of Chicago Institute for Neuroscience start-up funds, the NIH (1DP2NS111657-01) and (1RF1NS127123-01) awarded to M.S and a T32 training grant (T32DA043469) from National Institute on Drug Abuse awarded to S.K..

## Additional Information

### Contributions

Conceptualization: S.K. M.S. ; Data curation: C.D., S.K. ; Formal analysis: Y.C. ;, Investigation: Y.C., M.S., C.D., S.K. ; Visualization: Y.C. ; Writing - original draft : Y.C., M.S. ; Writing - review and editing: Y.C., M.S., C.D., S.K.

### Data availability

Raw imaging data are extremely large and not feasible for upload to an online repository but are available upon request.

---

**Figure 1—figure supplement 1.**
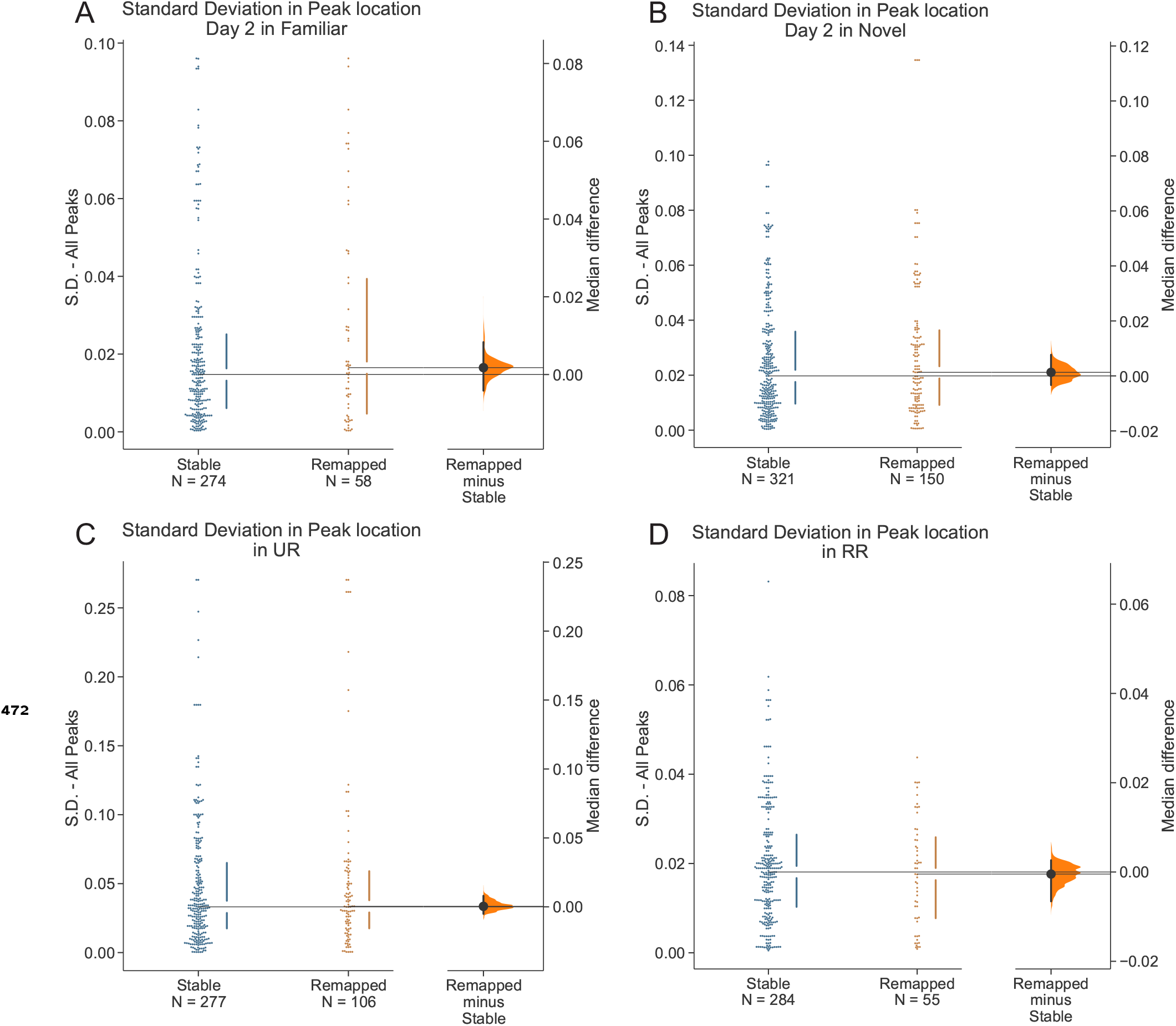
Following the initial episode, remapped and stable PFs have similar spatial precision during subsequent episodes. (A) Comparison of spatial precision of PFs on day 2 in the familiar environment. Right, bootstrapped median difference between remapped vs stable PFs from day 1. 5000 re-samples, P = 0.421. (B) Comparison of spatial precision of PFs on day 2 in the novel environment. Right, bootstrapped median difference between remapped vs stable PFs from day 1. 5000 re-samples, P = 0.811. (C) Comparison of spatial precision of PFs in UR condition. Right, bootstrapped median difference between remapped vs stable PFs from R. 5000 re-samples, P = 0.869 (D) Comparison of spatial precision of PFs in RR condition. Right, bootstrapped median difference between remapped vs stable PFs from R. 5000 re-samples, P = 0.848

**Figure S1—figure supplement 2.**
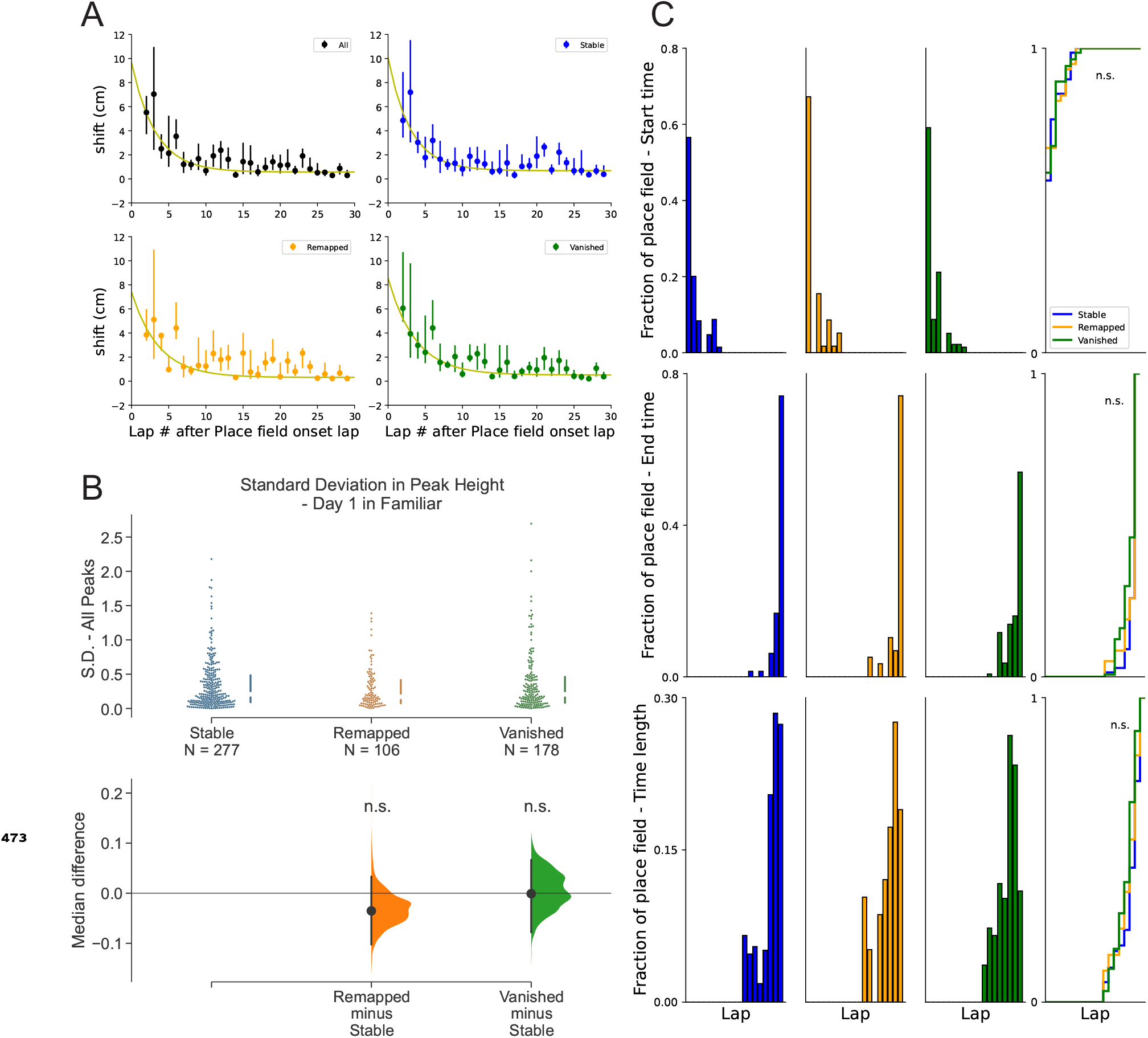
Other place field metrics are not associated with place field fate across days in a familiar environment. (A) Comparison of backward shifting dynamics between the different PF fate categories

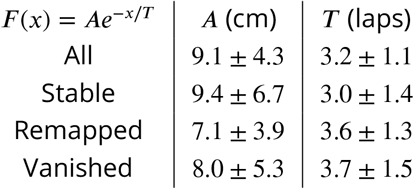

(B) Comparison of lap-by-lap variation in peak amplitude. P = 0.311 for stable vs remapped, 5000 re-samples, P = 0.993 for stable vs vanished(C) Histograms of PF onset laps, end laps, and duration (in laps) for Stable, Remapped and Vanished PFs. Cumulative fraction plots (right). Wilcoxon rank-sum test,

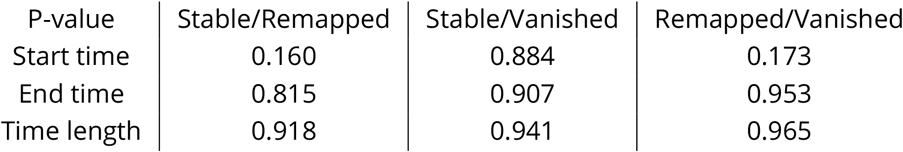

**Figure 2—figure supplement 1.**
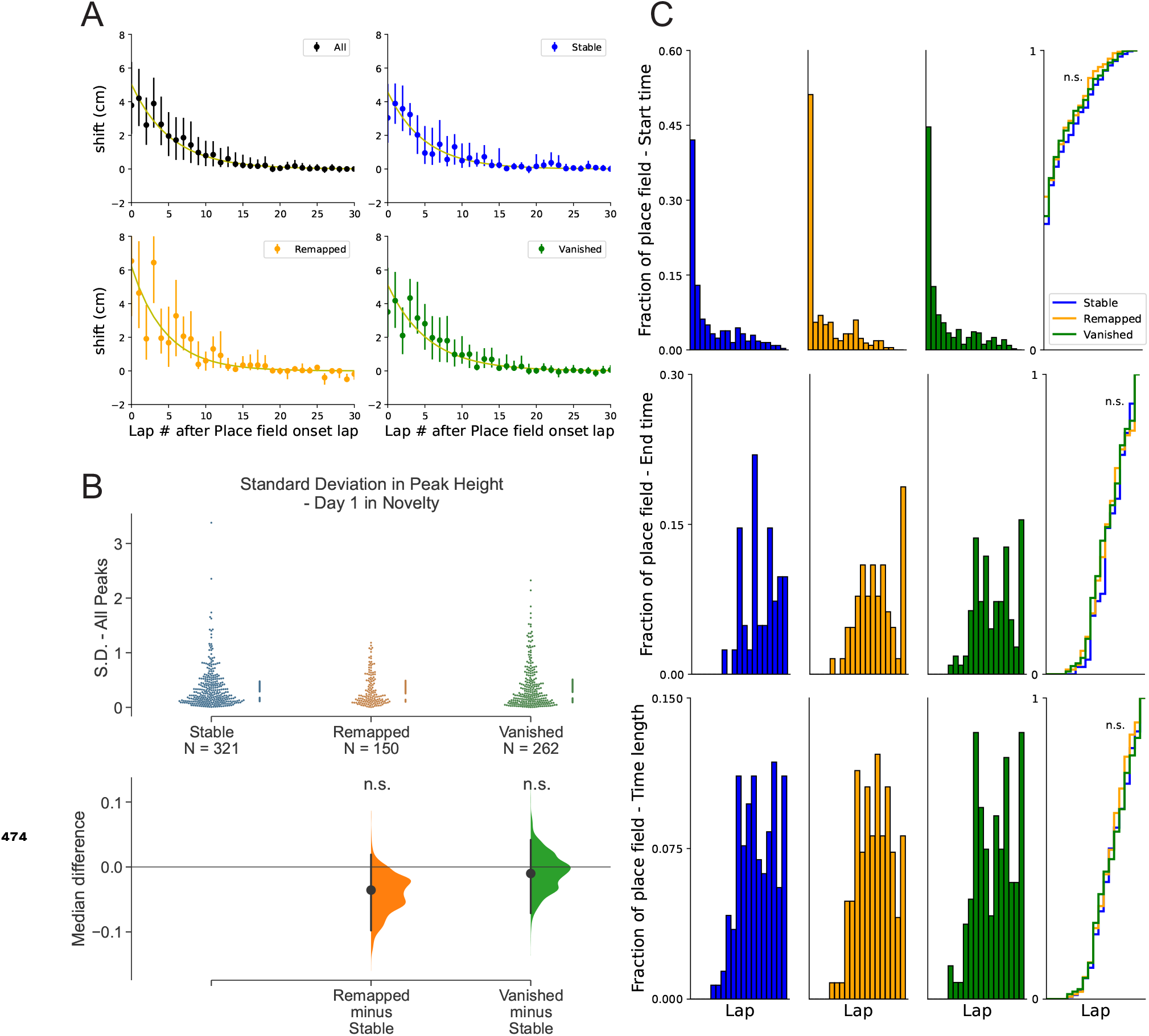
Other place field metrics are not associated with place field fate across days in a novel environment. (A) Comparison of backward shifting dynamics between the different categories of PF fate

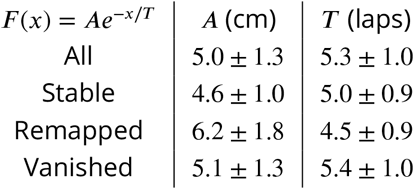

(B) Comparison of lap-by-lap variation in peak amplitude. P = 0.223 for stable vs remapped, 5000 resamples, P = 0.847 for stable vs vanished. (C) Histograms of PF onset laps, end laps, and duration (in laps) for stable, remapped and vanished PFs. Cumulative fraction plots (right). Wilcoxon rank-sum test,

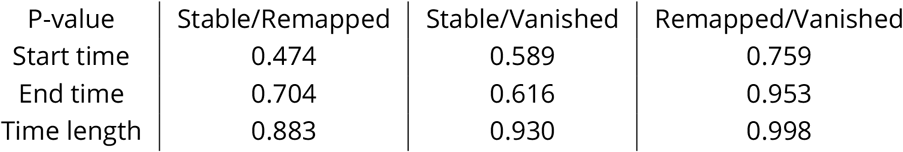

**Figure 3—figure supplement 1.**
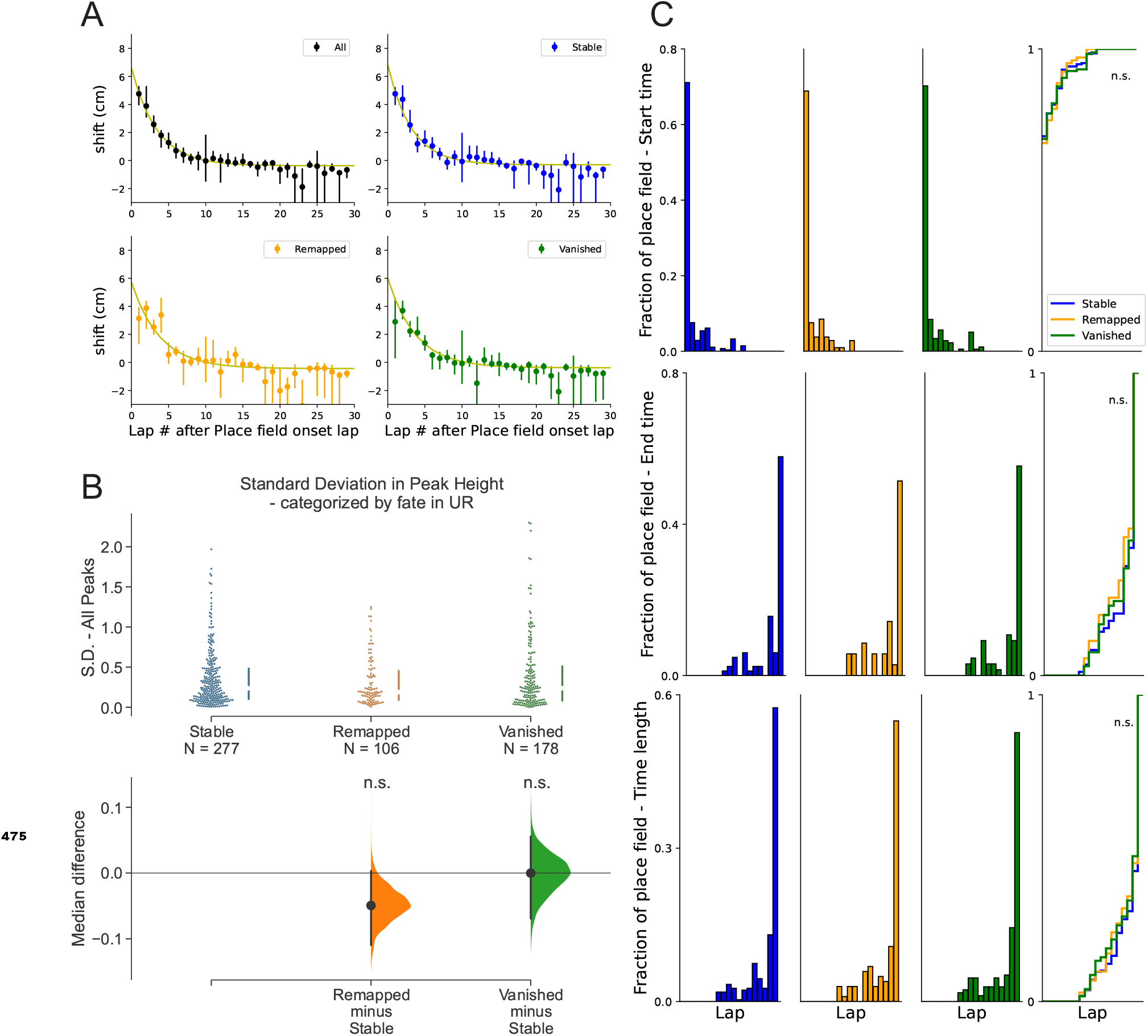
Other place field metrics in R are not associated with place field fate in UR. (A) Comparison of backward shifting dynamics between the different categories of PF fate

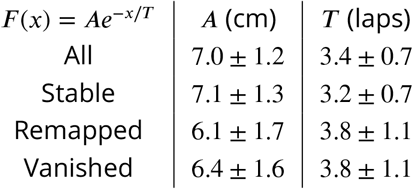

(B) Comparison of lap-by-lap peak amplitude variation. P = 0.140 for stable vs remapped, 5000 re-samples, P = 0.996 for stable vs vanished. (C) Histograms of PF onset laps, end laps, and duration (in laps) for stable, remapped and vanished PFs. Cumulative fraction plots (right). Wilcoxon rank-sum test,

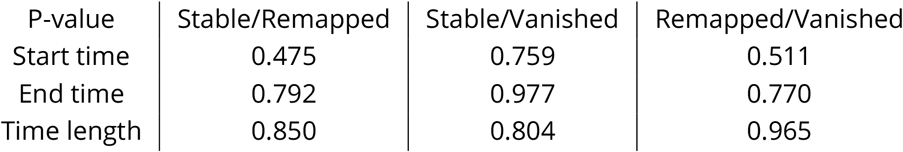

**Figure S3—figure supplement 2.**
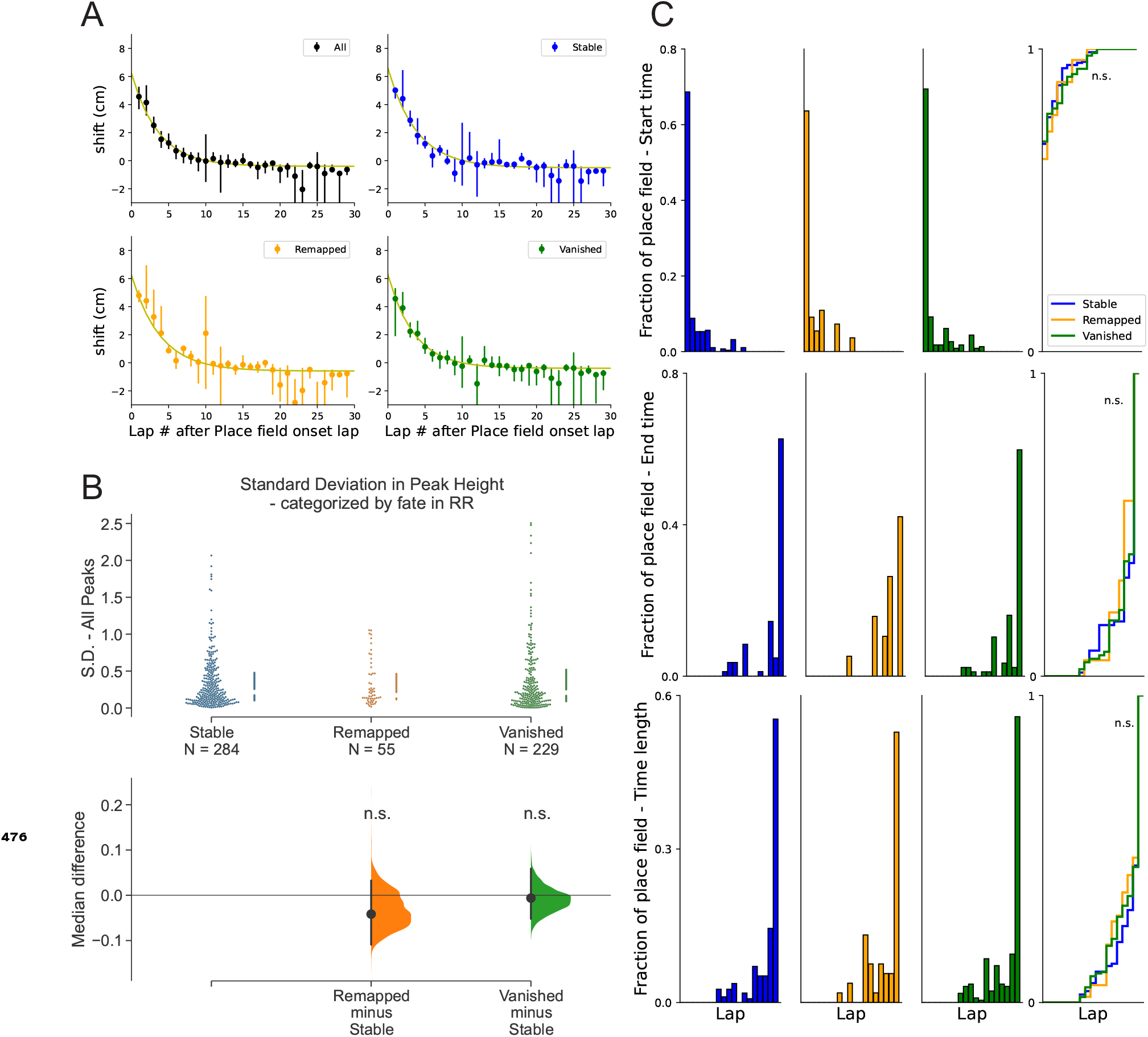
Other place field metrics in R are not associated with place field fate in RR. (A) Comparison of backward shifting dynamics between the different categories of PF fate

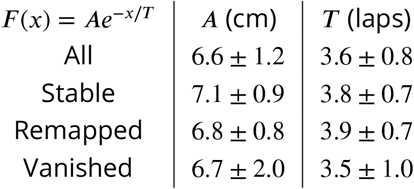

(B) Comparison lap-by-lap peak amplitude variation. P = 0.338 for stable vs remapped, 5000 re-samples, P = 0.680 for stable vs vanished. (C) Histograms of PF onset laps, end laps, and duration (in laps) for stable, remapped and vanished PFs. Cumulative fraction plots (right). Wilcoxon rank-sum test,

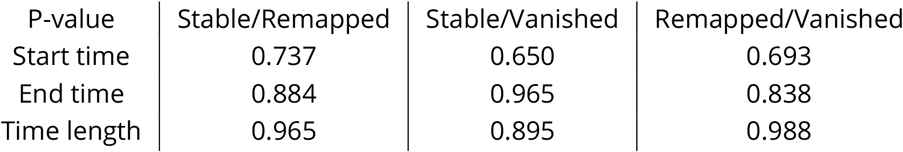

